# NAVIP: Unraveling the Influence of Neighboring Small Sequence Variants on Functional Impact Prediction

**DOI:** 10.1101/596718

**Authors:** Jan-Simon Baasner, Andreas Rempel, Dakota Howard, Boas Pucker

## Abstract

Once a suitable reference sequence has been generated, intraspecific variation is often assessed by re-sequencing. Variant calling processes can reveal all differences between strains, accessions, genotypes, or individuals. These variants can be enriched with predictions about their functional implications based on available structural annotations, i.e. gene models. Although these functional impact predictions on a per-variant basis are often accurate, some challenging cases require the simultaneous incorporation of multiple adjacent variants into this prediction process. Examples include neighboring variants which modify each other’s functional impact. The Neighborhood-Aware Variant Impact Predictor (NAVIP) considers all variants within a given protein coding sequence when predicting the effect. As a proof of concept, variants between the *Arabidopsis thaliana* accessions Columbia-0 and Niederzenz-1 were annotated. NAVIP is freely available on GitHub (https://github.com/bpucker/NAVIP) and accessible through a web server (https://pbb-tools.de).

**Author Summary:** Intraspecific variation gains increasing relevance as reference genome sequences are available for many investigated (plant) species. Understanding the effects of sequence variants between individuals of a population is a challenge. SnpEff (Cingolani et al., 2012) is the current standard tool for predicting the functional impact of sequence variants, but only considers one sequence variant at a time. We developed NAVIP to properly handle cases in which multiple sequence variants cluster together and influence each other’s functional impact. A comparison of two *Arabidopsis thaliana* accessions demonstrates the importance of considering multiple sequence variants simultaneously for the prediction of changes in encoded proteins. NAVIP is universally applicable to any organism for which the relevant sequence information and structural annotation is available. All underlying code is freely available on GitHub and we operate a web server for users’ convenience.

## Introduction

Re-sequencing projects examining many individuals or accessions of a species [1–4], are becoming increasingly important in plant research. Approaches similar to genome-wide association studies (GWAS) which are based on mapping-by-sequencing (MBS) are frequently applied in a wide range of crop species [5–8]. They are boosted by a rapidly increasing availability of high-quality reference genome sequences for crops [9–13], technological advances in long-read sequencing [14], and low sequencing costs [15,16]. *De novo* assemblies are still beneficial for the detection of large structural variants [17–22] and especially to reveal novel sequences [18,19,21,23], but the reliable detection of modifying single nucleotide variants (SNVs) can be achieved based on (short) read mappings. Well established tools for the small sequence variant discovery in plants are BMA MEM and GATK [24–27]. In recent years, long-read sequencing is gaining popularity in studies exploring the intraspecific diversity, as more sequence variants can be detected in previously inaccessible genomic regions [28,29]. One of the most frequently used tools for long read mapping is minimap2 [30] that can handle both relevant technologies, Pacific Biosciences and Oxford Nanopore Technologies, well. Hundreds of dedicated variant calling tools have been developed to harness the specific potential and to cope with challenges that come with long reads. Famous tools for the discovery of SNVs based on long reads are Longshot [31], SVIM-asm [32], and Sniffles2 [33]. One advantage of long reads is the ability to assign small sequence variants to different haplophases.

Once identified, the annotation of sequence variants is performed by predicting their functional implications based on the available gene models (structural annotation). Leading tools such as ANNOVAR [34], VEP [35], and SnpEff [36] currently perform this prediction efficiently by focusing on a single variant at a time. An impact prediction facilitates the identification of targets for post-GWAS analyses and can lead to the identification of small sequence variants that form the molecular basis of commercially relevant phenotypic differences [7,37,38]. Although the effect prediction for single variants is computationally efficient and usually correct, there are challenging cases in which predictions based on a single variant alone cannot be accurate. (1) Multiple InDels could either lead to frameshifts or they could compensate for each other’s effect leaving the sequence with minimal modifications [39–41] and (2) two SNVs occurring in the same codon could lead to a different amino acid substitution compared to the apparent effects resulting from an isolated analysis of each of these SNVs. It is important to note that SNVs and InDels can also influence each other’s effects.

Here we present a computational tool for accurately predicting the combined effect of phased variants on annotated coding sequences. The Neighborhood-Aware Variant Impact Predictor (NAVIP) was developed to investigate large variant data sets of plant re-sequencing projects, but is not limited to the annotation of variants in plants. As a proof of concept, NAVIP was used to identify cases between the *A. thaliana* accessions Columbia-0 (Col-0) and Niederzenz-1 (Nd-1) where an accurate impact prediction needs to consider multiple variants at a time.

## Results

### Features of NAVIP

NAVIP predicts the functional impact of sequence variants by considering all sequence variants affecting the coding sequence of a gene simultaneously. Users need to supply a set of sequence variants (VCF), a reference genome sequence (FASTA), and a structural annotation (GFF3). NAVIP returns an annotated VCF file and FASTA files with corrected coding and polypeptide sequences. If phased sequence variants are provided in the VCF file, NAVIP performs separate analyses for the different haplophases.

NAVIP can be retrieved from a GitHub repository (https://github.com/bpucker/NAVIP) and is executable without installation. Additionally, NAVIP is also available free of charge through a web server (https://pbb-tools.de/NAVIP). This makes NAVIP accessible to a wide range of users and applicable to data sets of various sizes. Uploaded files are used only for the intended analysis and are deleted 48 hours after offering the results for download. The web server is able to send notification emails upon completion of a job, which can serve as documentation and facilitate the analysis of large data sets.

### Relevance of NAVIP for prediction of premature stop codons

Running NAVIP on an *A. thaliana* Nd-1 data set with 644,261 SNVs (S1 File, S2 File) took about 5 minutes on a single core with a peak memory usage of about 3 GB RAM. To the best of our knowledge, SnpEff is the most frequently used tool for the annotation of variants and is also universally applicable. Therefore, the NAVIP output was compared with the SnpEff predictions generated for the same data set and structural annotation. The results are largely congruent, but interesting cases for comparison are predictions of premature stop codons, as these may have severe biological consequences. While a single SNV would cause a premature stop codon, the simultaneous presence of two SNVs can result in an amino acid encoding codon (**Figure 1a**). Of 600 premature stop codons predicted by SnpEff, 144 were identified as amino acid substitutions when considering multiple SNVs in the same codon via NAVIP (**Figure 1b**). Given the total of 600 predicted premature stop codons in this Nd-1 data set, 24% were false positive predictions. NAVIP revealed that tyrosine frequently occurs instead of a premature stop codon because the tyrosine codons are very similar to two of the three stop codons. There are also 17 additional premature stop codons predicted by NAVIP, which are the consequence of two sequence variants affecting the same codon. Despite the surprisingly large difference between the SnpEff and NAVIP results when it comes to predicting premature stop codons, the differences in affected genes are smaller. Many genes with a predicted premature stop codon have multiple downstream premature stop codons. While the prediction of an individual premature stop codon might be wrong for a certain position, the gene can still be correctly identified by both tools as harboring premature stop codons if additional ones occur further downstream (S3 File). If a premature stop codon results in a loss-of-function event, the accumulation of additional variants is likely due to a lack of purifying selection. To support the assumption that genes with premature stop codons lost their function, the rate of amino acid changing variants in these genes was compared to all other genes (**Figure 1c, Figure 1d**). The number of variants changing amino acids (aa_N_) to those resulting in the same amino acid (aa_S_) was calculated for all genes (aa_N_/aa_S_). A significantly higher proportion of amino acid changing variants was observed in genes with predicted premature stop codons compared to all other genes (Mann-Whitney U test, p-value=10^−161^). Premature stop codons might frequently appear in genes undergoing pseudogenization that are barely expressed, as purifying selection would be weak or even absent in these cases. Therefore, we investigated the expression of genes with premature stop codons in *A. thaliana*. A comparison of the average expression of genes with a premature stop codon against all other protein encoding genes (**Figure 1e, Figure 1f**) revealed a significantly lower expression of genes with premature stop codons (Mann-Whitney U test, p-value=10^−70^).

**Figure 1:**
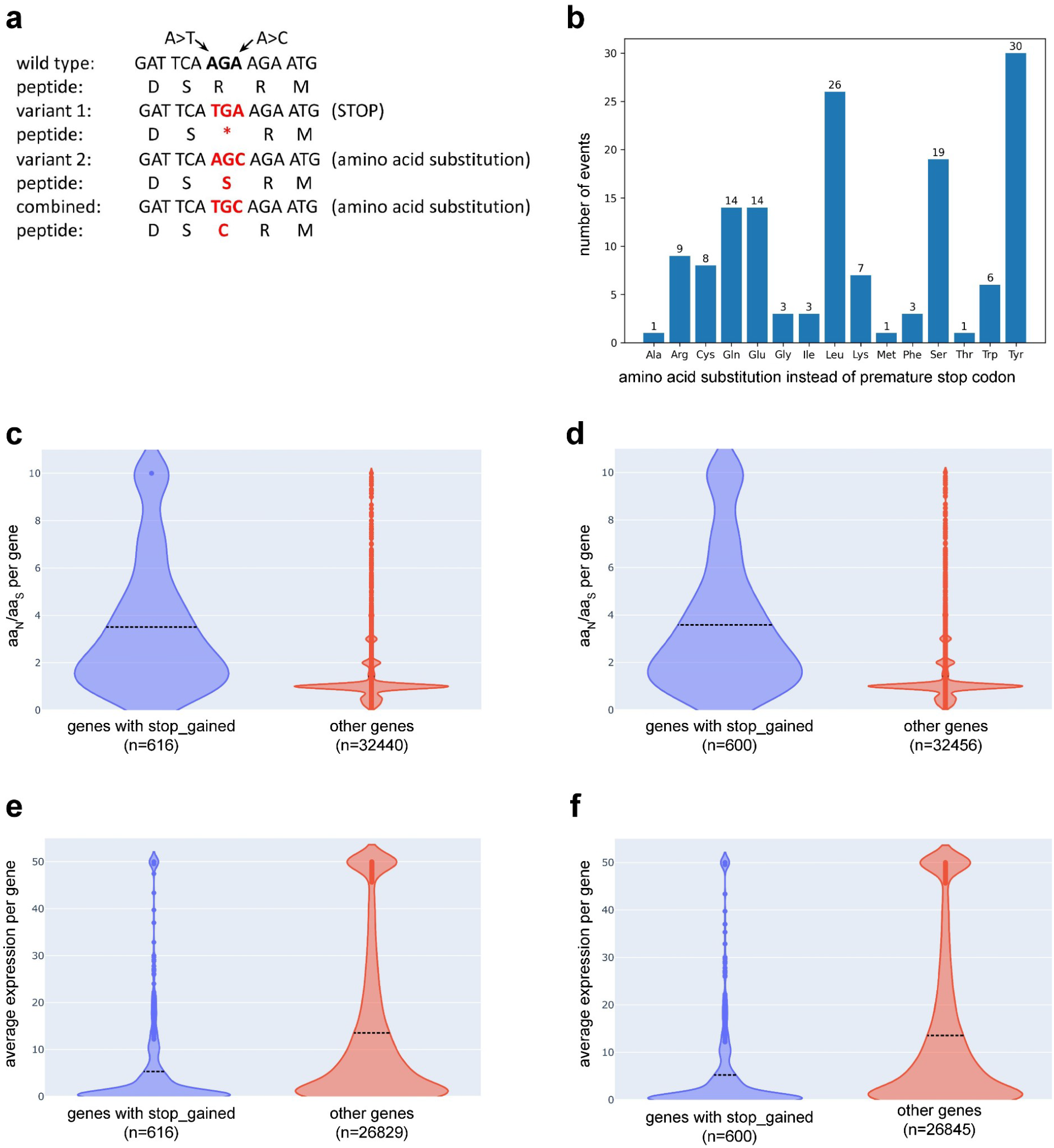
(a) This illustration shows the concept of two SNVs affecting the same codon resulting in different prediction outcomes. (b) Second site variants within the same codon turn premature stop codons predicted by SnpEff into amino acid substitutions. In 144 cases, premature stop codons are substitutions by the respective amino acids. (c) The proportion of amino acid changing variants is significantly higher in genes with premature stop codons predicted by NAVIP (blue) compared to all other genes (red). aaN is the number of variants changing an amino acid residue and aaS is the number of variants resulting in the same amino acid residue. (d) The proportion of amino acid changing variants is significantly higher in genes with premature stop codons predicted by SnpEff (blue) compared to all other genes (red). Data underlying these visualizations are available in S3 File. (e) Comparison of the average expression of genes with a premature stop codon predicted by NAVIP against all other protein encoding genes with available expression data. (f) Comparison of the average expression of genes with a premature stop codon predicted by SnpEff against all other protein encoding genes with available expression data.

To demonstrate the scalability of NAVIP, we processed 200 samples from the 1135 accession comparison study [1]. On average, an accession harbored 498 cases of stop codons predicted by SnpEff were classified as amino acid substitutions by NAVIP (S4 File).

While premature stop codons are probably the most severe changes, we also explored the influence of neighboring SNVs on amino acid substitutions between Col-0 and Nd-1. A total of 50,122 amino acid substitution predictions were analyzed including cases in which one of the annotation tools predicts no change of the amino acid. Predictions of NAVIP and SnpEff were congruent in 46,680 cases (93.1%) and differed in 3442 cases (6.9%) (S5 File).

### Role of compensating InDels (cInDels)

InDels can compensate for each others’ frameshift when occurring together in the same haplophase (**Figure 2a**). While the first InDel can alter the reading frame, the second one could revert the reading frame back to the original one, thus resulting in only a few altered codons enclosed by the two events. Since premature stop codons can emerge in the novel codons following the first frameshift, the distance between such InDels is expected to be very small. An analysis of the distance distribution of the InDels between Nd-1 and Col-0 (S6 File) revealed that most compensating InDels (cInDels) occur within a very short distance of 2-8 bp (**Figure 2b**). Multiples of three are more frequent than other distances of a similar size, which might be connected to the length of codons. Since *A. thaliana* is considered highly homozygous, we assume that all identified sequence variants are located in the same haplophase.

**Figure 2:**
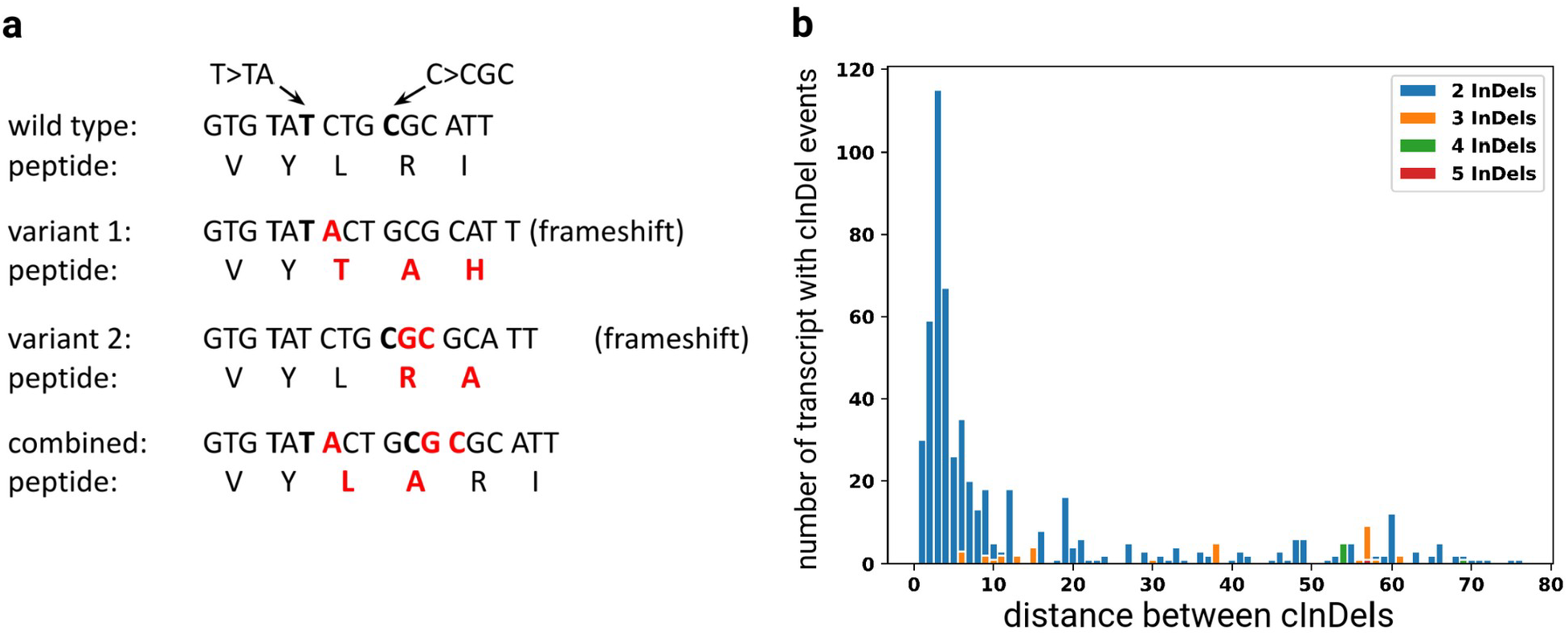
(a) Theoretical concept of two InDels compensating each others’ frameshift. The first insertion changes the reading frame, while the second insertion shifts the reading frame back to the original one. While each individual variant would suggest a loss-of-function due to a frameshift mutation, the combination of both results in ’only’ two additional amino acids in the gene product. (b) Distribution of distances between compensating InDels (cInDels). As the second InDel can compensate for the frameshift caused by the upstream InDel, distances between such cInDels are short and frequently multiples of three. In total, 484 genes were identified to contain cInDels in the Nd-1 data set.

## Discussion

This study demonstrates features of NAVIP by utilizing a previously generated set of high confidence sequence variants [26]. There is always a trade-off between sensitivity and specificity in the variant calling process [26,42] (see S1 File for details). The benchmarking of NAVIP is conducted by comparing it with SnpEff, which controls for the quality of the sequence variant dataset to minimize its impact on the results. As an additional validation of the outcome, NAVIP results were analyzed for additional amino acid substitutions in genes with premature stop codons. The frequency of such variants was higher in genes with premature stop codons compared to others suggesting a lack of purification selection in these genes which could point to pseudogenization. The comparison against all other genes also clearly revealed the increased frequency of amino acid substitutions in genes with premature stop codons. Additionally, a low expression of genes with premature stop codons compared to other genes suggests a pseudogenization. In summary, the properties observed for genes with premature stop codons match the expectations thus supporting the biological validity of the data set.

One motivation for the development of NAVIP was to fill a gap that exists between variant calling and variant annotation software. Variant calling involves the identification of genetic variants from raw sequencing data. This process typically features algorithms that analyze read alignments and uses statistical models to detect variants. Variant callers such as GATK [60] produce VCF files containing potential genetic variants. Variant annotation, on the other hand, assigns functional relevance to identified variants. This step requires databases and algorithms to provide additional information about each variant. Annotation tools such as ANNOVAR [34], VEP [35], or SnpEff [36] process VCF files previously generated by callers, rather than performing the variant calling themselves, thus losing access to the original read information. The separation between these two steps is due to technical and conceptual differences and serves several purposes. First, a separation of concerns: Variant calling focuses on the detection of variations, while annotation concentrates on the interpretation of those variants, allowing for specialized optimization of each step without complicating the other. Second, computational efficiency: Calling variants requires processing raw sequencing data, which can be computationally intensive. A streaming application would need to stop processing and accumulate all variants until there is complete gene information before annotating, which can be challenging in terms of memory usage, especially for large genes or when dealing with many samples simultaneously. Thus, separating the annotation step from the initial variant calling allows for a more efficient use of computational resources. Third, data flow and scalability: By separating calling and annotation, researchers can perform these steps independently, allowing for parallel processing and easier scaling of analysis pipelines. The VCF format used in variant calling is optimized for documenting detected variants, while other annotation formats are better suited for downstream analysis.

We developed NAVIP to simultaneously assess the impact of all neighboring sequence variants in protein encoding sequences and to be universally applicable. The described cases in the comparison of two *A. thaliana* accessions demonstrate the necessity to have such a tool at hand. NAVIP revealed the presence of second site mutations that compensate for other variants, e.g. turning a presumed premature stop codon into an amino acid substitution or vice versa. Another example are frameshifts resulting from InDels that are compensated by downstream InDels, which shift the reading frame back to the original pattern. Neglecting these interactions of sequence variants during the functional impact prediction can lead to mis-annotation. While NAVIP was developed to accurately predict changes in the polypeptide sequence based on DNA sequence variants, downstream tools are needed to predict consequences of these changes on the function of proteins. Tools like SIFT [43], PolyPhen-2 [44], or SNAP2 [45] could be applied for this next step. Many computational tools for the assessment of DNA sequence variant impact focus on human data sets [46–49]. The objective is often to identify pathogenic variants [43,50]. Universally applicable tools like SnpEff [36], which are also suitable to analyze plant data sets, predict the impact of isolated sequence variants. The purpose of NAVIP is to offer novel functionalities to the plant science community and other communities working on non-model organisms. NAVIP could boost the power of re-sequencing studies by opening up the field of compensating or in general mutually influencing variants. Such variants have the potential to reveal new insights into patterns of molecular evolution and especially co-evolution of sites. The consideration of multiple variants during the effect prediction could reveal novel targets in GWAS-like approaches. The availability through a web server enables a large community of scientists without computational skills to benefit from NAVIP.

The remaining challenge is now the reliable detection of sequence variants prior to the application of NAVIP. A range of tools is available for the mapping of short reads and the following identification of sequence variants [26]. There is also rapid progress in the development of long read mapping tools [51,52] and the subsequent variant identification [53– 56]. For heterozygous and polyploid species, phasing of these variants is another task that needs to be addressed in the future. Variant callers could directly report multiple SNVs of one haplophase as one MNV by collapsing the individual variants. In contrast to variant callers, variant annotators do not have access to the aligned reads and cannot infer this information. The correct prediction of functional implications relies on the correct assignment of variants to respective haplophases. If provided with accurately phased variants, NAVIP can perform predictions for highly heterozygous and even polyploid species. Previous studies demonstrated that sequence variants might only affect individual isoforms in a negative way [50]. NAVIP analyzes all annotated transcript isoforms and would be able to discover such cases. Currently, a major limitation is the lack of isoform-resolved annotation for non-model plant species. Given the rapid progress in long read sequencing [14,57,58], it is likely that highly accurate structural annotation will become available for most plant species in the next few years.

## Materials and Methods

### Implementation of the Neighborhood-Aware Variant Impact Predictor (NAVIP)

The Neighborhood-Aware Variant Impact Predictor (NAVIP) (https://github.com/bpucker/NAVIP) has been implemented in Python3. NAVIP requires a VCF file containing sequence variants, a FASTA file containing the reference sequence, and a GFF3 file containing the structural annotation (gene models) as input. The variants provided must be homozygous or in a phased state to allow an accurate impact prediction per allele. If no information about the phasing is provided, all variants are assumed to be in the same haplophase. Effects on all annotated transcripts are evaluated per gene by taking into account the presence of all given variants simultaneously. NAVIP consists of three modules: VCF preprocessing, the NAVIP main program, and a simple first analysis (SFA) of the generated annotation. The first module is designed to preprocess VCF files line-by-line to check for multiallelic variants, i.e. variants with more than one alternative allele at a given position, split them into two separate entries, and convert them into one of three categories: substitution, insertion, or deletion. This process is crucial, as it allows for a clearer representation, facilitating further analysis and interpretation. The preprocessing also removes conflicting data entries and logs warnings and potential errors, such as identical bases, to ensure that any encountered discrepancies are documented for review. The second module is designed to validate genetic variants against transcript sequences, with a particular focus on insertions and deletions, to ensure that the variants align correctly with the reference and match the corresponding sequences in the transcript. NAVIP generates a new VCF file with an additional annotation field and additional report files. One annotation string in the VCF output file matches the SnpEff result format, but also has a NAVIP-specific string with additional information (see the manual for details: https://github.com/bpucker/NAVIP/wiki). NAVIP also produces FASTA files with sequences harboring all variants. NAVIP enhances the VCF files by incorporating additional information about the variants, including their effects on coding sequences (CDS), codon changes, and amino acid alterations. This allows users to identify variants with a potential impact on protein function, providing researchers with deeper insight into the effects of genetic variation. Frameshift mutations can occur when the number of nucleotides inserted or deleted is not a multiple of three, altering the downstream amino acid sequence. The third module serves as a primary interface for identifying compensating insertions and deletions (cInDels) within a given VCF file, categorizing them based on their effect on the reading frame, and generating output files summarizing the findings. It also includes functionality to visualize the number of InDels across transcripts through bar plots, facilitating interpretation of the results. The automatic assessment of complementing InDels reveals the relevance of simultaneously considering all InDels within a coding sequence when predicting their impact. All NAVIP scripts can be downloaded from the above-mentioned GitHub repository and do not require the installation of any dependencies other than the Python packages. NAVIP is also available through a web server (https://pbb-tools.de/NAVIP) free of charge. Files are kept confidential and will be deleted 48 h after offering the results for download.

### Identification and validation of sequence variants

Illumina sequencing reads of *A. thaliana* Nd-1 [17] were mapped to the *A. thaliana* Col-0 reference genome sequence (TAIR10) [59] via BWA MEM v.0.7.13 [24] using the –m option to avoid spurious hits. Variant calling was performed via GATK v3.8 [60] based on the developers’ recommendation. This combination of BWA MEM and GATK was previously identified as a reliable approach for this particular data set [26]. All processes were wrapped into Python scripts (https://github.com/bpucker/variant_calling) to facilitate automatic execution on a high-performance compute cluster. An initial variant set was generated based on hard filtering criteria recommended by the GATK developers. The two following variant calling runs considered the set of surviving variants from the previous round as the gold standard to avoid the need for hard filtering.

Since a high-quality genome sequence assembly of Nd-1 was previously generated [18], we harnessed this sequence to validate all variants identified by short-read mapping. From the start of each chromosome sequence, variants sorted by genomic position were successively tested by taking the upstream sequence from Col-0, modifying it according to all upstream *bona fide* variants, and searching for it in the Nd-1 assembly (S7 File). Variants were admitted to the following analysis if the assembly supported them. This consecutive inspection of all variants enabled a reliable removal of false positives, leading to a set of high-confidence variants. The genome-wide distribution of the sequence variants was assessed using a previously developed Python script [17].

An independent confirmation of randomly selected sequence variants was performed using Sanger sequencing. *A. thaliana* Nd-1 plants were grown as previously described [17] to extract DNA from leaf tissue using a cetyltrimethylammonium bromide (CTAB)-based method [61]. Oligonucleotides flanking the regions that harbor the variants of interest were designed manually (S8 File). Amplification via PCR, analysis of PCR products via agarose gel electrophoresis, purification of PCR products, Sanger sequencing, and evaluation of results were following previously established protocols [62].

### Comparison of NAVIP and SnpEff stop gain prediction

To the best of our knowledge, SnpEff [36] is the most widely used tool for predicting the effects of sequence variants, thus it was selected for comparison. NAVIP can only provide more accurate effect predictions if multiple sequence variants interfere, e.g. if multiple SNVs are located within the same codon. Otherwise, the predictions of NAVIP and SnpEff would be the same. Consequently, the following comparison focuses only on cases of multiple sequence variants that might interfere with each other.

SnpEff v4.1f [36] was applied with default parameters to the *A. thaliana* Nd-1 variant data set to predict the effects of SNVs based on the Araport11 [63] structural annotation of the TAIR10 genome sequence of *A. thaliana* Columbia-0. NAVIP was also applied to the same data set for benchmarking. Predictions of premature stop codons were compared between NAVIP and SnpEff results, as these cases have the potential to show biologically important differences. This analysis was performed exclusively on SNVs to avoid the influence of frameshifts that would be caused by InDels. Only the most upstream predicted premature stop codon within any gene was considered in this analysis. To support the loss of function of the affected genes, the frequency of amino acid changing variants (aa_N_) was compared to the number of variants that did not alter the encoded amino acid (aa_S_). This ratio was compared between genes with premature stop codons and all other genes, expecting a higher ratio of variants that change the encoded amino acids if the gene undergoes pseudogenization. The Python package plotly was used to visualize these data distributions in violin plots. A pseudocount was added to both aa_N_ and aa_S_ to enable the ratio calculation in case when aa_S_ would be 0. aa_N_/aa_S_ ratios greater than 10 were set to this maximum value to enable visualization. A Mann-Withney U test was performed using Python to test for significant differences between the two groups. When genes with a premature stop codon undergo pseudogenization, they may show lower than average gene expression. Therefore, a comparison of the expression of genes with a premature stop codon against all other protein-coding genes was performed. A previously compiled count table based on all publicly available paired-end RNA-seq data sets of *A. thaliana* [64] was harnessed for this analysis. Differences were visualized using the Python package plotly as described above, with the expression values clipped at 50 to enable an informative visualization. All Python scripts developed for these analyses are freely available on GitHub (https://github.com/bpucker/variant_calling).

### Assessment of compensating InDels (cInDels)

An independent analysis of insertions/deletions (InDels) was performed by NAVIP to understand the relevance of considering all InDels within a CDS simultaneously. Transcripts with predicted frameshifts were analyzed to identify downstream insertions/deletions which are compensating each other’s effect, i.e. the second frameshift is reverting an upstream frameshift. The distance between these events was analyzed by NAVIP and is included in the standard output. This analysis is not restricted to pairs of cInDels, but can also handle multiple InDels compensating each other’s frameshifts.

## Availability and requirements

Project name: NAVIP

Project homepage: https://github.com/bpucker/NAVIP

Operating system(s): Linux (website is platform independent)

Programming language: Python3

Other requirements: Python3

License: GNU General Public License v3.0

RRID: SCR_024838

## Data availability

The data sets supporting the results of this article are publicly available or included within the article and its additional files. Python scripts developed and applied for this study are available on GitHub: https://github.com/bpucker/NAVIP (https://doi.org/10.5281/zenodo.10613052) and https://github.com/bpucker/variant_calling (https://doi.org/10.5281/zenodo.10613055).

## Declarations

### Authors’ contributions

BP designed the research. JSB wrote the NAVIP code. AR updated the NAVIP code and added NAVIP to the pbb-tools web server. JSB, DH, and BP conducted bioinformatics analyses. DH and BP performed the experimental validation. JSB, AR, and BP wrote the manuscript. All authors read and approved the final version of the manuscript and agreed to its submission.

### Financial Disclosure Statement

The authors received no specific funding for this work.

### Competing Interests

JSB, AR, and DH have no competing interests. BP is head of the technology transfer center Plant Genomics and Applied Bioinformatics at iTUBS. This does not alter our adherence to PLOS policies on sharing data and materials.

## Acknowledgments

We acknowledge support by members of Genetics and Genomics of Plants, Bioinformatics Resource Facility, and Sequencing Core Facility at the Center of Biotechnology. We thank Hanna Schilbert for critical reading of the manuscript. We thank the Center for Biotechnology (CeBiTec) at Bielefeld University for providing an environment to perform the computational analyses. Many thanks to the German network for bioinformatics infrastructure (de.NBI, grant 031A533A) and the Bioinformatics Resource Facility (BRF) at the Center for Biotechnology (CeBiTec) at Bielefeld University for providing an environment to perform the computational analyses. We acknowledge support by the Open Access Publication Funds of Technische Universität Braunschweig.

## Supporting Information

**S1 File**: Detailed description of the variant calling process, the validation process, and the resulting sequence variant data set.

**S2 File**: VCF file containing SNVs between Nd-1 and Col-0.

**S3 File**: Detailed information about premature stop codons predicted by NAVIP and/or SnpEff.

**S4 File**: Differences in the effect prediction between SnpEff and NAVIP for 200 accessions of the 1,135 *Arabidopsis thaliana* accession resequencing project.

**S5 File**: Comparison of SnpEff and NAVIP prediction differences between Col-0 and Nd-1. The table lists matches and differences for each possible amino acid substitution type.

**S6 File**: VCF file containing InDels between Nd-1 and Col-0.

**S7 File**: Schematic illustration of the variant validation process.

**S8 File**: FASTA file containing oligonucleotide sequences used for the generation and sequencing of amplicons to validate randomly selected sequence variants.

## Notes

https://github.com/bpucker/NAVIP

https://github.com/bpucker/variant_calling

### Summary of Updates

- analysis of some of the 1135 A. thaliana accession re-sequencing data added - detailed description of NAVIPs functions

