## Supplementary material for "NAVIP: Unraveling the Influence of Neighboring Small Sequence Variants on Functional Impact Prediction": S1 File

#### Variant detection and validation

Nd-1 reads were mapped against the Col-0 reference genome sequence (TAIR10). Based on 124,662,140 mapped paired-end reads, 384,622 variants were detected in the first variant calling round. This initial set was extended over three additional rounds of variant calling leading to over one million variants. The variant calling was stopped, because no substantial increase in the number of novel variants was observed during the last rounds. An assembly based on independent Single Molecule Real Time (SMRT) sequencing reads supported 772,643 (76.6%) of all variants detected during the last iteration (Figure 1, Additional file 4, Additional file 6). On average, one variant was observed every 154 bp between Col-0 and Nd-1. SNV frequencies ranged from one event in 225 bp on Chr5 to one event in 158 bp on Chr4. InDel frequencies ranged from one event in 1,051 bp on Chr5 to one event in 809 bp on Chr4.

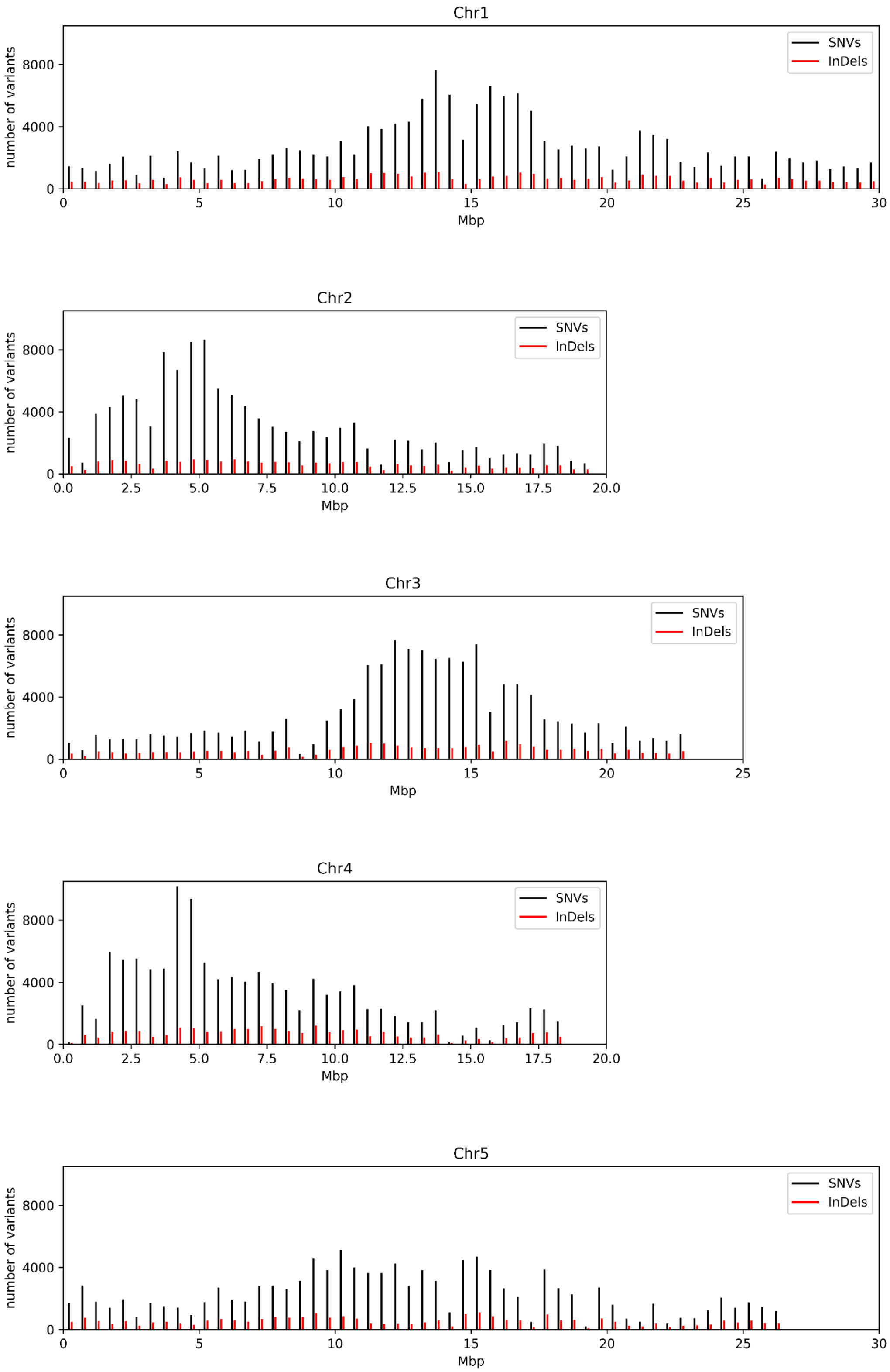


**Figure 1**: Genome-wide distribution of sequence variants between Col-0 and Nd-1. Distributions of SNVs and InDels over the chromosome sequences of Col-0 were visualized as previously described (Pucker *et al.*, 2016).

Although the repeated variant calling processes were intended to increase the sensitivity, we did not observe a substantial improvement between the second and third round. This saturation indicates that no additional variants would be detected in further variant calling rounds. The number of detected variants as well as the validation rate was almost constant (Table 1).

**Table 1**: Numbers of total and validated variants.

| Variant data set | Total variants | Validated variants |
|---|---|---|
| **Initial set based on hard filtering** | 384,617 | 350,005 (90.1%) |
| **Soft filtering round 1** | 1,006,920 | 771,449 (76.6%) |
| **Soft filtering round 2** | 1,008,610 | 772,612 (76.6%) |
| **Soft filtering round 3** | 1,008,629 | 772,643 (76.6%) |

#### Experimental validation

Randomly selected loci with two SNVs within one codon were experimentally validated via PCR followed by amplicon sequencing (Table 2). Successful sequencing reactions show a validation rate of >95%.

**Table 2**: Neighboring SNVs validated in Nd-1 via PCR and amplicon sequencing. Sequences of oligonucleotides used for the amplicon generation are listed in Additional file 2.

| AGI | Fw primer | Rv primer | Status |
|---|---|---|---|
| At1g30545 | N400 | N401 | Validated |
| At3g55500 | N402 | N403 | Validated |
| At3g26770 | N406 | N407 | Validated |
| At4g30570 | N408 | N409 | Validated |
| At1g28150 | N410 | N411 | Validated |
| At1g35430 | N412 | N413 | Validated |
| At4g27230 | N414 | N415 | One SNV failed |
| At5g60230 | N424 | N425 | 2 validated |
| At1g31820 | N426 | N427 | 4 validated / 1 failed |

### Discussion of the variant identification and validation

Although differentiation between *bona fide* variants (true positives) and false positives based on a high quality genome sequence assembly worked very well, false negatives were not taken into account and might even bias this classification approach by preventing the validation of neighboring variants (Additional file 1). If a variant is missed by the initial variant calling, its presence in the flanking sequence used during the validation process will prevent a proper match. Therefore, the number of variants could be slightly higher than reported here. Nevertheless, this conservative approach was selected to minimize the risk of keeping false positive variants. There is always a trade-off between sensitivity and specificity in the variant calling process (Olson *et al.*, 2015) and our approach is in strong favor of specificity. However, the number of identified and validated variants exceeds previous reports of 485,887 variants between Col-0 and Nd-1 (Pucker *et al.*, 2016). Instead the observed variant frequency is closer to the results of a comparison between Bur-0 and Col-0 (Ossowski *et al.*, 2008). Despite the difference in total numbers, the distribution on the chromosome scale is similar to the previous comparison of Col-0 and Nd-1 (Pucker *et al.*, 2016). It seems that Chr4 is the most variable one, while Chr5 is the least variable one between both compared accessions. Successful validation via PCR and amplicon sequencing supported the presence of two SNVs within one codon. Although these variants are perceived as two SNVs, the underlying mechanism could be a multiple nucleotide polymorphism (MNP). It would be interesting to see if these SNVs occur independently in other accessions in the *A. thaliana* population.
