## Supplementary figures and images for "NAVIP: Unraveling the Influence of Neighboring Small Sequence Variants on Functional Impact Prediction"

### S7 File

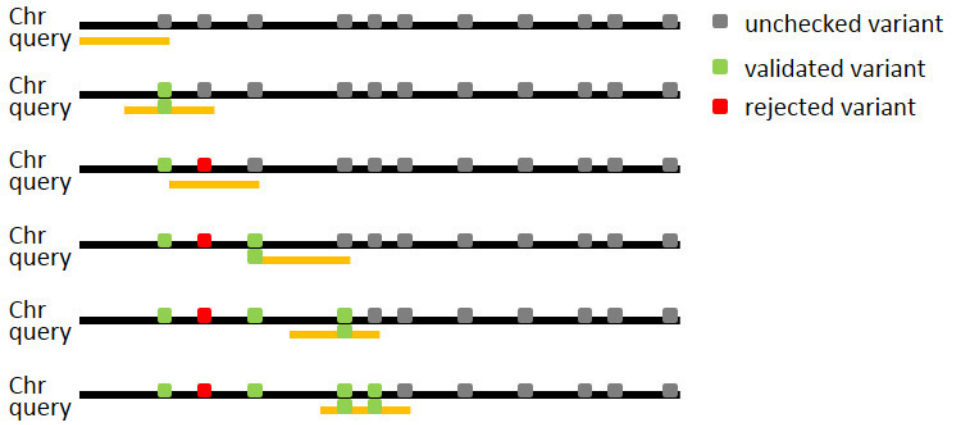
